# Angiogenesis in the mature mouse cortex is governed in a region specific and Notch1 dependent manner

**DOI:** 10.1101/2024.05.24.595778

**Authors:** Alejandra Raudales, Ben Schager, Dominique Hancock, Sorabh S. Sharma, Kamal Narayana, Patrick Reeson, Manjinder Cheema, Jakob Körbelin, Craig E. Brown

**Affiliations:** Division of Medical Sciences, University of Victoria, Victoria, BC, Canada; Department of Oncology, Hematology and Bone Marrow Transplantation, University Medical Center Hamburg-Eppendorf, Germany; Department of Psychiatry, University of British Columbia, Vancouver, BC, Canada

**Keywords:** angiogenesis, microcirculation, plasticity, Notch, VEGF, rarefaction, capillary, blood flow

## Abstract

Cerebral angiogenesis is well appreciated in development and after injury, but the extent to which it occurs across cortical regions in normal adult mice and underlying mechanisms, is incompletely understood. Using *in vivo* imaging, we show that angiogenesis in anterior-medial cortical regions (retrosplenial and sensorimotor cortex), was exceptionally rare. By contrast, angiogenesis was significantly elevated in posterior-lateral regions such as visual cortex, primarily within 200µm of the cortical surface. There were no regional differences in vessel pruning or sex effects except for the length and depth of new capillaries. To understand mechanisms, we surveyed gene expression and found Notch related genes were enriched in ultra-stable retrosplenial versus visual cortex. Using endothelial specific knockdown of Notch1, cerebral angiogenesis was significantly increased along with genes implicated in angiogenesis (*Apln, Angpt2, Cdkn1a*). Our study shows that angiogenesis is regionally dependent and manipulations of Notch1 signaling could unlock the angiogenic potential of the mature vasculature.

---

The brain uses up to 20% of the energy metabolised by the body to sustain the needs of neurons and glial cells^1^. Despite the tremendous efficiency of this system, there is little capacity for energy storage in the form of glycogen, therefore the brain requires a constant supply of blood through the cerebrovascular system. It follows that alterations or interruptions to blood flow, even within the smallest vessels known as capillaries, can have deleterious effects on cognitive and sensori-motor function^2^. There is a substantial body of literature showing the vascular system can rapidly adapt (in a matter of seconds to minutes) to meet the brains’ needs, largely through modulating vessel tone^3–5^. Over longer time scales, the vascular system can eliminate or generate new blood vessels from existing ones, referred to as pruning and angiogenesis, respectively. This form of structural plasticity is well appreciated in brain development^6–9^, where there is abundant vascular endothelial cell growth along gradients of hypoxia, to support nascent brain regions. Mechanistically, new vessel growth in development is primarily orchestrated by vascular endothelial growth factor (VEGF) receptor signalling pathways in growing tip cells^10–13^. VEGF receptors in tip cells interact with NOTCH signalling, which play a critical role in vessel development by stabilizing endothelial stalk cells, thereby allowing lumen formation and patency of new vessels^14,15^. However, mechanistic studies in adulthood typically involve hypoxia or stroke to induce angiogenic activity, therefore whether similar signalling pathways are required for ongoing angiogenesis in the healthy mature brain, is not well understood.

Although there is considerable evidence showing that mature vascular networks remodel when challenged by hypoxic stimuli, stroke, diabetes or exercise^16–27^, the extent to which these networks can change under normal, healthy conditions is debatable. Aside from histological studies that assume a constitutive but low level of angiogenesis in the healthy cortex^28–30^, *in vivo* 2-photon imaging studies have found little or no ongoing angiogenesis in the adult mouse cortex, even after weeks of voluntary exercise^6,31–35^. However, virtually all studies focus on one or occasionally two cortical regions, typically somatosensory and/or motor cortex. Thus, there has yet to be a systematic survey of vascular remodelling across cortical regions.

This possibility of regional differences in vascular plasticity has recently come into focus with multiple reports showing brain region specific differences in susceptibility to capillary plugging, neurovascular coupling, and vascular density^36–40^. Indeed, work from our lab has revealed that some cortical regions such as the visual cortex, are resistant to age related vessel loss, whereas others (ie. retrosplenial, somatosensory/motor cortex) are not^38^. This raises the possibility that there may be inherent differences in the rate at which cortical regions produce or eliminate vessels. In order to address this question, we used longitudinal two-photon imaging in adult mice to survey vascular remodelling across multiple cortical regions that span the anterior to posterior extent of dorsal cortex. Our results reveal a surprising level of angiogenic potential in visual cortex, whereas retrosplenial, forelimb/hindlimb somatosensory or motor cortex exhibited very little. Further we link regional differences in angiogenesis with *Notch1* gene expression, which appears to serve as a brake on cerebral angiogenesis given that endothelial knockdown significantly enhanced angiogenesis.

## Results

### Patterns of blood vessel growth in mature cortex

To examine the sprouting of new blood vessels or their elimination (angiogenesis vs. pruning) in the adult mouse cortex, we implanted a cranial window over anterior or posterior cortical regions in 2–3 month old female and male C57BL6/J mice (**Fig. 1a, b**, n=15 and 7 mice, respectively). About 4 weeks later, we employed intrinsic optical signal (IOS) imaging to functionally map the visual and hindlimb somatosensory cortex for posterior windows (see Vis and HL in **Fig. 1b, c**), or hindlimb and forelimb somatosensory cortex for anterior windows. Mapping these areas provided landmarks to delineate other cortical regions such as retrosplenial, motor, and trunk/whisker somatosensory cortex (RS, Mo and Tr/W in **Fig. 1b**). Ten days after IOS mapping, we used 2-photon microscopy to image vascular networks labelled by injection of 70kDa FITC dextran (i.v. 3-5% dissolved in saline) in each cortical region over a 23 day period (**Fig. 1d**). As shown in **Figure 1d**, each region could be reliably imaged to a depth of 400µm below the pial surface. Further, the vast majority of vessels could be re-identified for structural analysis indicating that: a) vascular networks are largely stable in adulthood and b) our surgical and imaging preparation induced minimal damage or hypoxia^41^.

**Figure 1.**
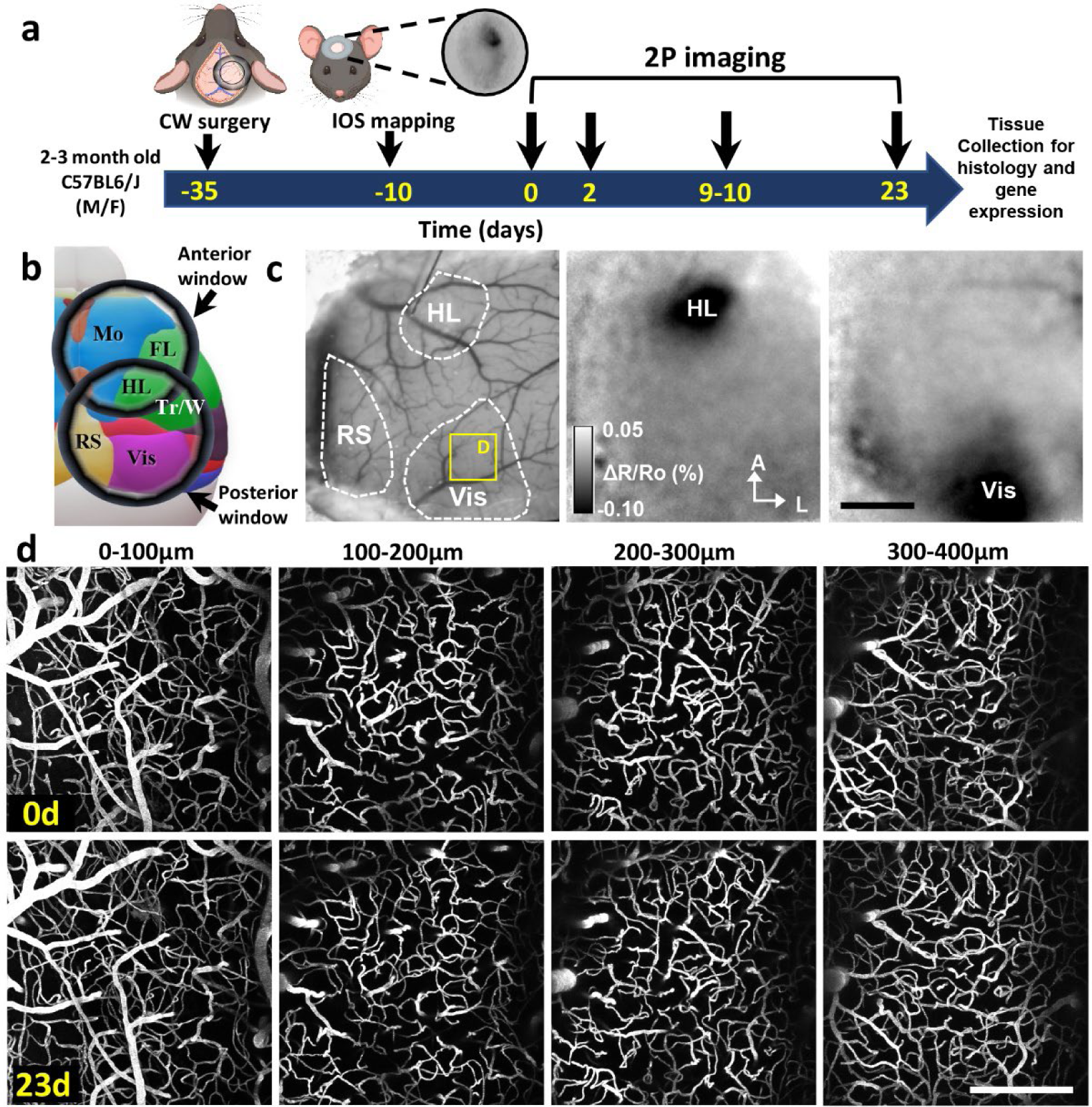
Longitudinal *in vivo* imaging of vascular remodelling across cortical regions in adult mice. (**a**) Timeline of experimental procedures and two-photon (2P) imaging sessions. (**b**) Schematic showing different cortical regions imaged with anterior or posterior positioned cranial windows. (**c**) Left: Brightfield image of the cortical surface showing the location of retrosplenial (RS), hindlimb somatosensory (HL) and Visual cortex (Vis). Middle and right: intrinsic optical signal (IOS) maps show the position of the hindlimb and visual cortex. (**d**) Maximal intensity z-projection images of fluorescently labelled blood vessels at different depths in the visual cortex at the start of the experiment and then again 23 days later. Mo: motor cortex, FL: forelimb somatosensory cortex, Tr/W: trunk and whisker-barrel somatosensory cortex. Scale bars = 1mm (c) and 200µm (d).

Although mature vascular networks are highly stable, we clearly observed instances of capillary sprouting and pruning over the 23 day imaging period (mean diameter 3.78±0.86µm, **Fig. 2a**). Since capillaries were imaged by fluorescent labelling of blood plasma, angiogenic sprouts often appeared to show a “blob or head-like” bumps at the growing end of the vessel (**Fig. 2a and 2d**), consistent with previous *in vivo* descriptions^20,33^. Our own post-mortem immunolabelling experiments in young mice confirmed that growing vessels often show a blob-like intravascular dye distribution (**Supp. Fig. 1a**). In general, the emergence or growth of new blood vessels followed one of three patterns. We therefore defined an angiogenic event as the formation of a new, connected blood vessel at day 23 that emerged from either a sprout or no sprout on day 0 (see examples in **Fig. 2b,c**). We also included examples where a sprout appeared at day 23 (at least 10µm in length), but was not yet connected to another vessel by the end of the experiment (**Supp. Fig. 1b**). In order to describe the progression of angiogenesis, we first identified all new blood vessels or sprouts at day 23, and then examined earlier time-points (day 0 and 9) to see how these new vessels/sprouts came about. On day 0, the majority of new vessels/sprouts (evident on day 23) had yet to show any signs of sprouting from the parent branch (“absent” 80%), while about 20% displayed a sprout (**Fig. 2d**). By day 9-10, 60% of vessels were either still in the sprouting stage or were already connected to another vessel, while 40% still had yet to show signs of a sprout. By day 23, 90% of all new vessels were connected to another vessel (see blue and green bars in **Fig. 2d**), with most showing evidence of blood flow (ie. streaking pattern in vessel lumen created by movement of red blood cells), while the remaining 10% were still in the sprout phase. To visualize how these angiogenic events progress, we plotted individual events at each phase of growth in **Figure 2e** (Note purple lines are events that end in a sprout, teal lines are those ending in a connected capillary). With respect to where newly formed or pruned vessels occur, we examined capillary branch orders from the nearest penetrating arteriole or ascending venule (see PA or AV in **Fig. 2f**). Our analysis agrees with previous early development work^9,42^, in that almost all angiogenic events occurred off of lower order capillary branches (branches 2-5) from the ascending venule rather than penetrating arteriole (**Fig. 2g**). Similarly for vessel pruning, most events occur off of branches of the ascending venule (**Fig. 2g**). These experiments show that new capillaries primarily originate from the ascending venule, which can occur within 9-10 days time but the majority (90%) take longer and become complete (and flowing) by day 23.

**Figure 2.**
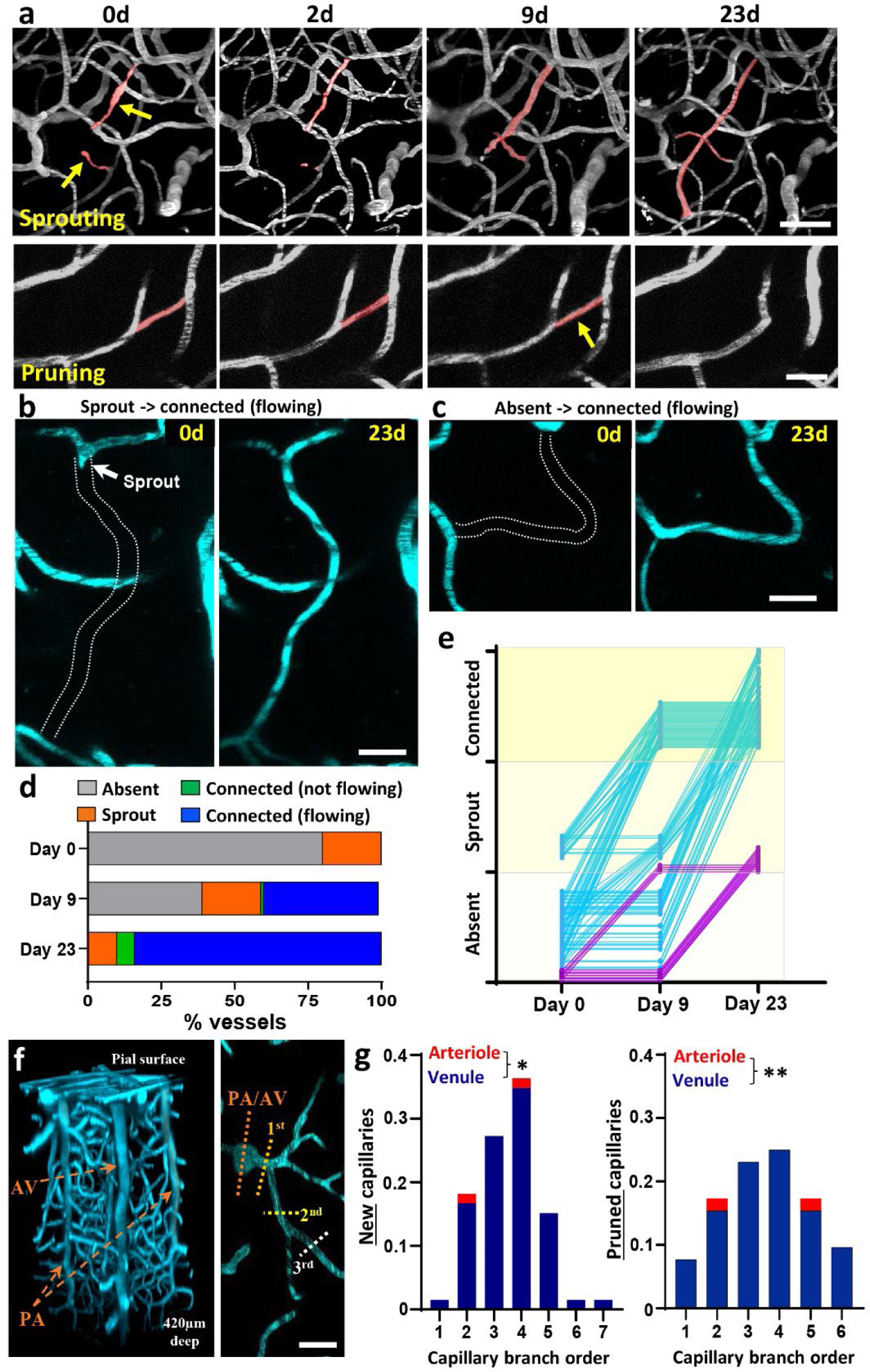
Progressive formation of new capillaries originate from branches off the ascending venule. (**a**) Time lapse *in vivo* maximal intensity z-projection images show development of a newly formed (top row) or pruned (bottom row) capillaries over 23 days in visual cortex. Newly formed capillaries with blood flow on day 23 originated from a sprout (**b**) or no sprout (“absent”, **c**) on day 0. (**d**) Graph shows time-dependent changes in the fraction (%) of newly formed capillaries at each stage of development (n=100 events). (**e**) Time course of each angiogenic event across the 3 phases of growth. Purple lines indicate events that finish day 23 as a sprout whereas teal lines indicate events that finish as a connected new capillary. (**f**) Left: maximal intensity y-z image projection showing the vasculature in visual cortex from the pial surface to 400µm deep. Right: branch order of new or pruned capillaries was determined by tracing back to the nearest penetrating arteriole (PA) or ascending venule (AV). (**g**) Almost the entire fraction of new (left side, 64/66 capillaries) or pruned capillaries (right side, 50/52 capillaries) originated from lower order capillary branches off the ascending venule. Data in g analysed with two-tailed Mann-Whitney test. *p≤0.05, **p≤0.01. Scale bars = 40µm (a, top row) and 20µm (a, bottom row, b, c, f).

### Cerebral angiogenesis but not pruning, is regionally dependent

To help reconcile discrepancies in the literature about rates of angiogenesis in the mature cortex, we imaged vascular networks in 6 different regions across the dorsal cortical mantle (RS, Vis, Tr/W, HL, FL and Mo, in general we could image 2-4 brain regions per mouse). Our analysis indicated a highly significant effect of brain region on the number of angiogenic events per mm^3^ (**Fig. 3a**; F_(5,64)_=5.80, p<0.001). As shown in **Figure 3a**, the highest rates were found in visual cortex whereas the lowest levels were in retrosplenial cortex. Contrasting with regional gradients in angiogenesis, vessel pruning did not vary significantly across cortical regions (**Fig. 3b**; F_(5,65)_=0.70, p=0.63). Summing up angiogenic and pruning events to provide an estimate of vessel turnover (or remodelling) also revealed a significant effect of brain region (**Fig. 3c**; F_(5,65)_=2.80, p<0.05), where turnover was highest in visual cortex and lowest in retrosplenial cortex. While no regional differences in pruning were found, linear regression indicated that regions with higher levels of angiogenesis also tended to have higher pruning rates (**Fig. 3d**, 69 areas imaged, R^2^=0.196, p<0.001). In order to better appreciate spatial gradients across the cortex, we plotted the stereotaxic coordinates (relative to bregma) of every imaging stack and expressed normalized rates of angiogenesis and pruning. The resultant map reveals a distinctive gradient in angiogenesis which was lower in anterior and medial regions and higher in posterior-lateral regions (**Fig. 3e**). These findings reveal that rates of cerebral angiogenesis, but not pruning are regionally dependent.

**Figure 3.**
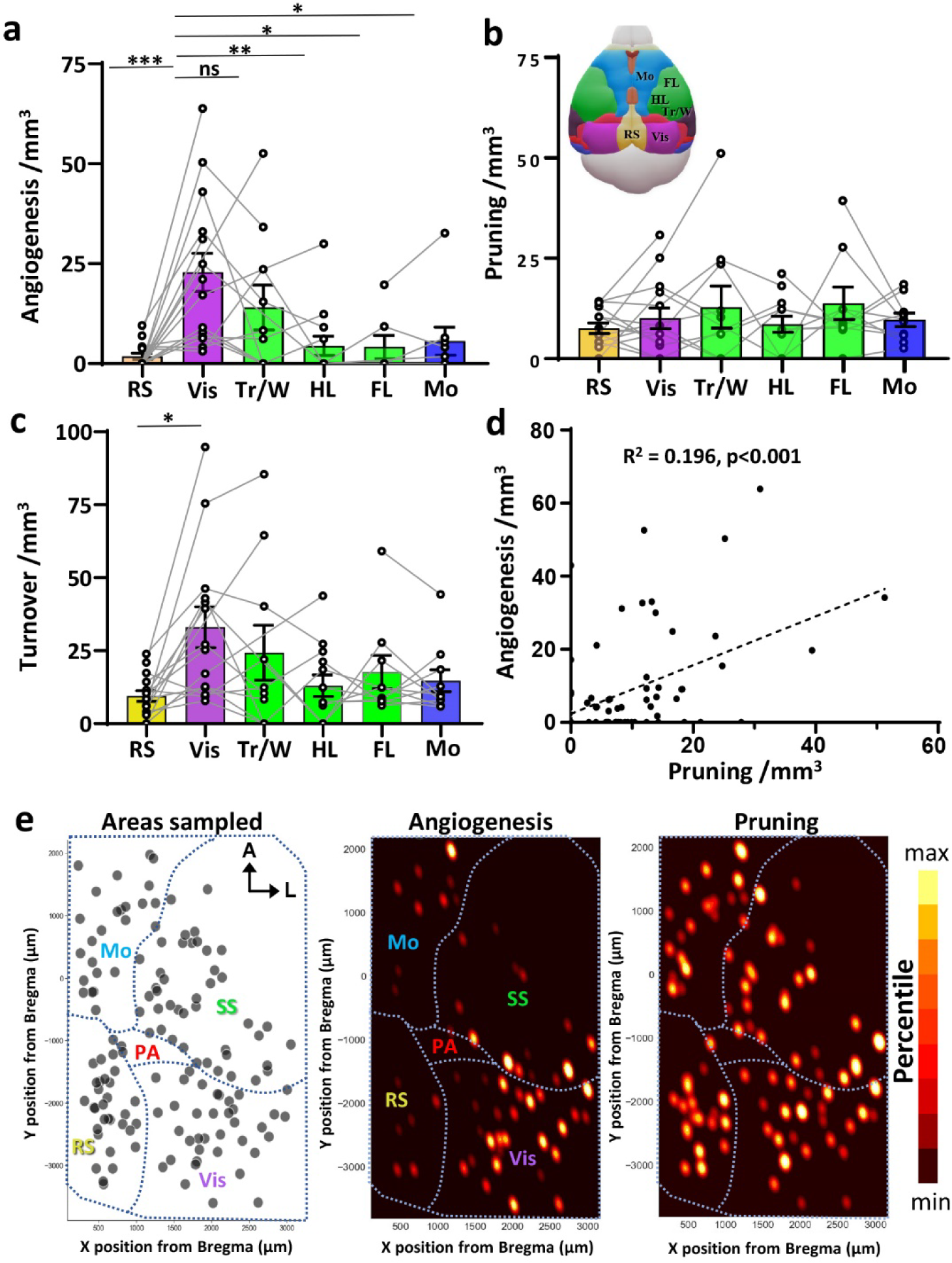
Cerebral angiogenesis but not pruning, is regionally dependent. (**a**) Graphs show individual data points (regions imaged in the same mouse are connected by lines) and the average number of angiogenic capillaries per mm^3^ across different cortical regions (sexes pooled together; n=16, 15, 10, 13, 7 and 9 mice for the 6 regions, respectively). (**b**) Individual and mean rates of capillary pruning per mm^3^ across different cortical regions. (**c**) Bar graph shows the sum of new and pruned capillaries (“turnover”) per mm^3^ in each cortical region. (**d**) Linear regression shows significant relationship between the mean rate of angiogenesis and pruning for each region imaged (n=68 regions). (**e**) Maps show the spatial location of each imaging stack across the cortical mantle relative to bregma (139 stacks from 24 mice) and corresponding weighted probability density of angiogenesis and pruning events for each stack. Color scale is proportional density for each plot, with maximum to minimum values expressed in 10^th^ percentile increments. Total area under all densities sums to 1. Data in a, b, c were analysed with one-way ANOVA followed by Tukey’s multiple comparisons tests. ns = not significant, *p≤0.05, **p≤0.01, ***p≤0.001. Data are mean ± SEM.

Regional differences in angiogenic rates could conceivably be related to: a) their relative position in the cranial window (ie. regions closer to window edge may be exposed to more damage during surgery) or b) differing proportions of ascending venules vs. penetrating arterioles (AV/PA), since new capillaries mainly originate from the venular side. Regression analysis indicated there was no relationship between rates of angiogenesis and a region’s distance to the nearest edge of the cranial window (r^2^=0.0005, p=0.8746; **Supp. Fig. 2a**). With regard to the latter issue, we sampled 21 imaging regions (n=7 V1; n=7 S1; n=7 RS) from 7 mice and identified 121 ascending venules or penetrating arterioles (AV/PA). In congruence with other studies^43^, we found a higher proportion of AVs (73/121; ∼60.3%) compared to PAs (48/121; ∼39.6%) across all regions. However, there were no differences in the proportions of AV vs. PAs across posterior cortical regions (**Supp. Fig. 2b**; AVs as % total: 60.0%, 58.5%, and 62.2% in RS, Visual and SS respectively).

Given there are some reports indicating that biological sex can influence cerebrovascular structure and function^44^, we stratified our experimental data accordingly. Since each sex was sampled similarly for mice with posterior placed cranial windows, we focused our analysis on retrosplenial, visual and somatosensory cortex that pooled HL and Tr/W areas (n=7 male and 8 female mice, respectively). As expected for angiogenic events (**Fig. 4a**), there was a highly significant effect of brain region (F_(2,35)_=8.40, p≤0.001) but no main effect of sex (F_(1,35)_=2.99, p=0.09) or sex by region interaction (F_(2,35)_=2.01, p=0.15). Analysis of pruning rates (**Fig. 4b**) did not show any differences based on region or sex (Main effect of sex: F_(1,35)_=2.28, p=0.14). We next examined whether sex influenced the depth where angiogenesis or pruning events occurred or the length of remodelled capillaries. For newly formed capillaries, these were deeper (**Fig. 4c**; t_(158)_=3.78, p<0.001) and longer in female mice (**Fig. 4d**; t_(158)_=4.62, p<0.0001) relative to their male counterparts. However, there were no sex dependent differences in the depth or length of pruned capillaries (**Fig. 4c, d**). Of note, pruned capillaries in both male and female mice were significantly shorter than angiogenic or stable capillaries (**Fig. 4d**). In summary, our data indicate that sex does not influence rates of cerebral angiogenesis or pruning but does affect the depth and length of newly formed capillaries.

**Figure 4.**
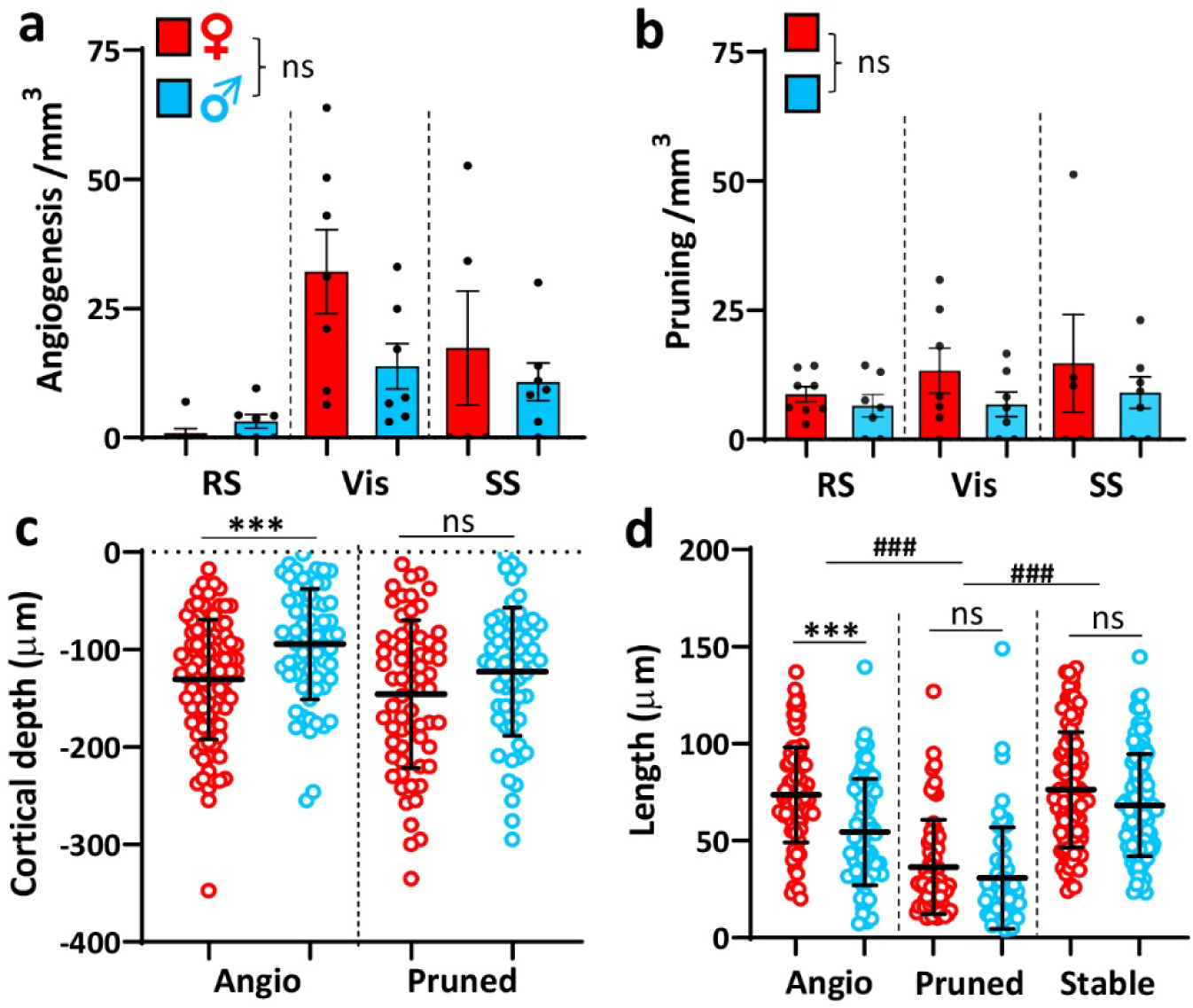
Sex affects the depth and length of newly formed capillaries but not rates of angiogenesis and pruning. (**a**) Bar graph shows individual and mean rate of angiogenesis per mm^3^ across different cortical regions in female and male mice (n=8 and 7 mice, respectively). SS represents pooled data from HL and Tr/W somatosensory regions. (**b**) Individual and mean pruning rates across cortical regions in male and female mice. (**c,d**) Graphs show the depth below the pial surface (**c**) and length (**d**) for each newly formed or pruned capillary in male and female mice (93 new capillaries in females vs 66 in males; 65 pruned capillaries in females vs 60 in males). The length of stable capillaries was based on a random sample of 100 capillaries from 4 mice per sex. Data in a and b were analysed with two-way ANOVA while data in c and d were analysed with unpaired two-tailed t-tests. ns = not significant, ***p≤0.001 for comparisons between sex. ### p≤0.001 for comparisons with sexes pooled. Data are mean ± SEM.

### Brain endothelial specific knockdown of Notch1 stimulates cortical angiogenesis

In order to understand the molecular mechanisms that regulate angiogenesis in the adult cortex, we surveyed gene expression of select signalling pathways previously implicated in vascular remodelling^45,46^. We focused our qPCR analysis on tissue from retrosplenial and visual cortex because they are situated relatively close to one another yet show the starkest differences in angiogenesis. These experiments revealed that genes associated with Notch (*Notch1*, *Jag1 and Dll4*) and VEGF receptor signalling (*Vegfr1, Vegfr2*) were more highly expressed in retrosplenial cortex relative to visual cortex (**Fig. 5a**). Since Notch ligands Dll4 and Jag1 signal through Notch1 receptors, which are expressed in endothelial cells and are more abundant in brain than other Notch receptors (Notch2-4)^47^, we examined whether endothelial knockdown of Notch1 signalling could stimulate cerebral angiogenesis *in vivo*. As shown in **Figure 5b**, we imaged the retrosplenial and visual cortex in homozygous *Notch1* floxed mice after saline injection (baseline) and then again after intravenous injection of AAV-BR1-iCre for endothelial specific knockdown of *Notch1*. To confirm Cre recombinase activity in endothelial cells, we injected cre-dependent Tdtomato reporter mice (Ai9) with saline and then again with AAV-BR1-iCre. Consistent with previous studies using this virus^48^, we found widespread and specific expression of cre-dependent reporter expression in cerebrovascular endothelial cells (**Fig. 5c, left** and **Supp. Fig. 3**; 84-89% of endothelial cells express reporter). As a control, we also ran a cohort of mice where AAV-BR1-GFP was injected first, followed several weeks later by AAV-BR1-iCre. Doing so prevents transfection of endothelial cells with AAV-BR1-iCre (**Fig. 5c, right**), likely due to the production of neutralizing antibodies associated with the first AVV injection. Analysis of our imaging data indicated that endothelial knockdown of Notch1 significantly increased rates of angiogenesis in both retrosplenial and visual cortex relative to baseline (see Sal/Cre in **Fig. 5d**). By contrast, our control group that were injected with AAV-BR1-GFP first, followed by AAV-BR1-iCre, did not show any changes in angiogenesis (see GFP/Cre in **Fig. 5d**). Rates of vessel pruning in both retrosplenial and visual cortex were not significantly affected by these manipulations (**Fig. 5e**). As an additional control, we injected mice with AAV-BR1-GFP alone and found no significant changes in angiogenesis or pruning when compared to the saline injected mice (**Supp. Fig. 4**). For the sake of completeness, we also examined the impact of endothelial specific knockdown of *Vegfr2* (injection of AAV-BR1-iCre in homozygous *Vegfr2* floxed mice), using the same experimental paradigm described above. This experiment did not reveal any significant effects of *Vegfr2* knockdown on rates of angiogenesis or pruning in retrosplenial and visual cortex (**Supp. Fig. 5**). Collectively, these results show that inhibiting Notch1 signalling in endothelial cells stimulates cortical angiogenesis, even in regions where angiogenesis is usually extremely limited (ie. retrosplenial cortex).

**Figure 5.**
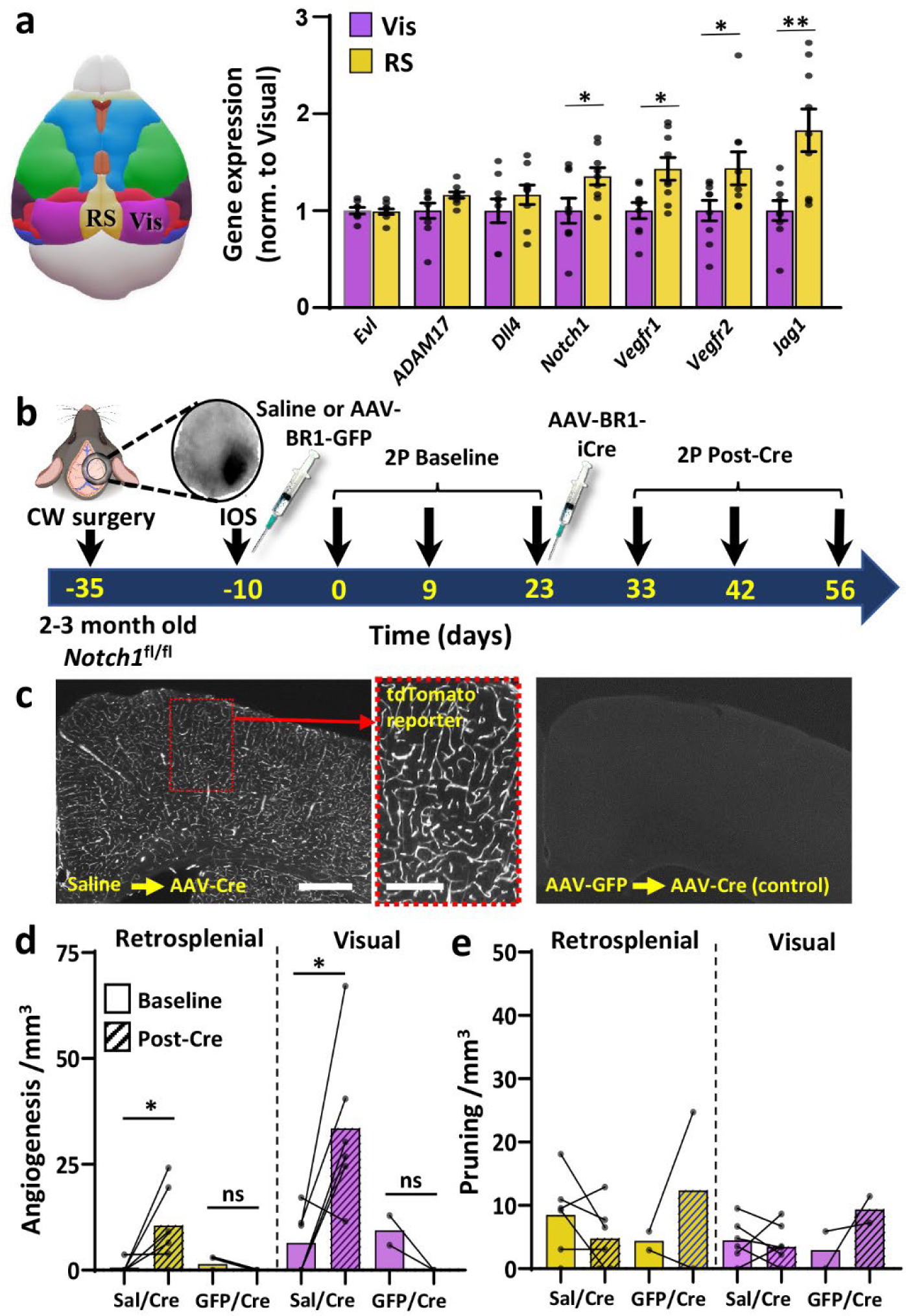
Endothelial knockdown of Notch1 signalling stimulates cerebral angiogenesis. (**a**) qPCR analysis shows gene expression from tissues collected from the retrosplenial and visual cortex (n=9 mice). Data are mean ± SEM. (**b**) Timeline of experimental procedures for imaging vasculature in retrosplenial and visual cortex before and after AAV mediated knockdown of endothelial Notch1. (**c**) Fluorescent images of coronal brain sections show cre-dependent expression of tdTomato in cortical vascular endothelial cells after injection of AAV-BR1-iCre (left) into Ai9 reporter mice. Right image: Ai9 mouse injected with AAV-BR1-eGFP at baseline, which do not show cre-dependent reporter expression after injection of AAV-BR1-iCre several weeks later. (**d**) Graphs show individual and mean rate of angiogenesis in retrosplenial and visual cortex in mice at baseline and then after with injection of AAV-BR1-iCre. Note that angiogenesis increases in mice with endothelial specific knockdown of Notch1 (Sal/Cre mice, n=6 mice), but not in control mice that were first injected with AAV-BR1-GFP at baseline (GFP/Cre, n=2 mice). (**e**) Pruning rates in retrosplenial and visual cortex in mice with endothelial specific knockdown of Notch1 or controls (Sal/Cre vs GFP/Cre mice). Data in a were analysed with unpaired two-tailed t-tests, while data in d, e were analysed with paired t-tests. ns = not significant, *p≤0.05, **p≤0.01. Scale bars in C = 0.5mm (left) and 0.2mm (middle).

In order to probe what genetic programs were induced in brain endothelial cells following *Notch1* knockdown, we performed bulk RNA sequencing on purified brain endothelial cells isolated from Notch1^flox/flox^ mice treated with AAV-BR1-iCre or GFP. Gene ontology (GO) analysis of differentially expressed genes (DEGs) revealed that Notch1 knockdown strongly altered expression of gene families implicated in angiogenesis (**Fig. 6a** and **Supp. Fig. 6a**). KEGG enrichment analysis of DEGs (up or down regulated) showed that Notch1 knockdown most significantly altered PI3K-AKT and MAPK signalling pathways (**Fig. 6b** and **Supp. Fig. 6b**). Quantification of differential gene expression indicated that Sy*t15, Tet1, Synpo2, Frem2* and *Notch1* were among the most significantly down-regulated genes, thus further validating our viral knockdown approach (**Fig. 6c,d**). Of note, several of the most significantly upregulated genes following Notch1 knockdown, namely *Apln, Angpt2, Cdkn1a, Plaur, Serpine1* to name a few, have been previously implicated in regulating angiogenesis (**Fig. 6c,d**). These findings indicate that Notch1 knockdown initiates a host of gene signalling pathways critical for angiogenesis.

**Figure 6.**
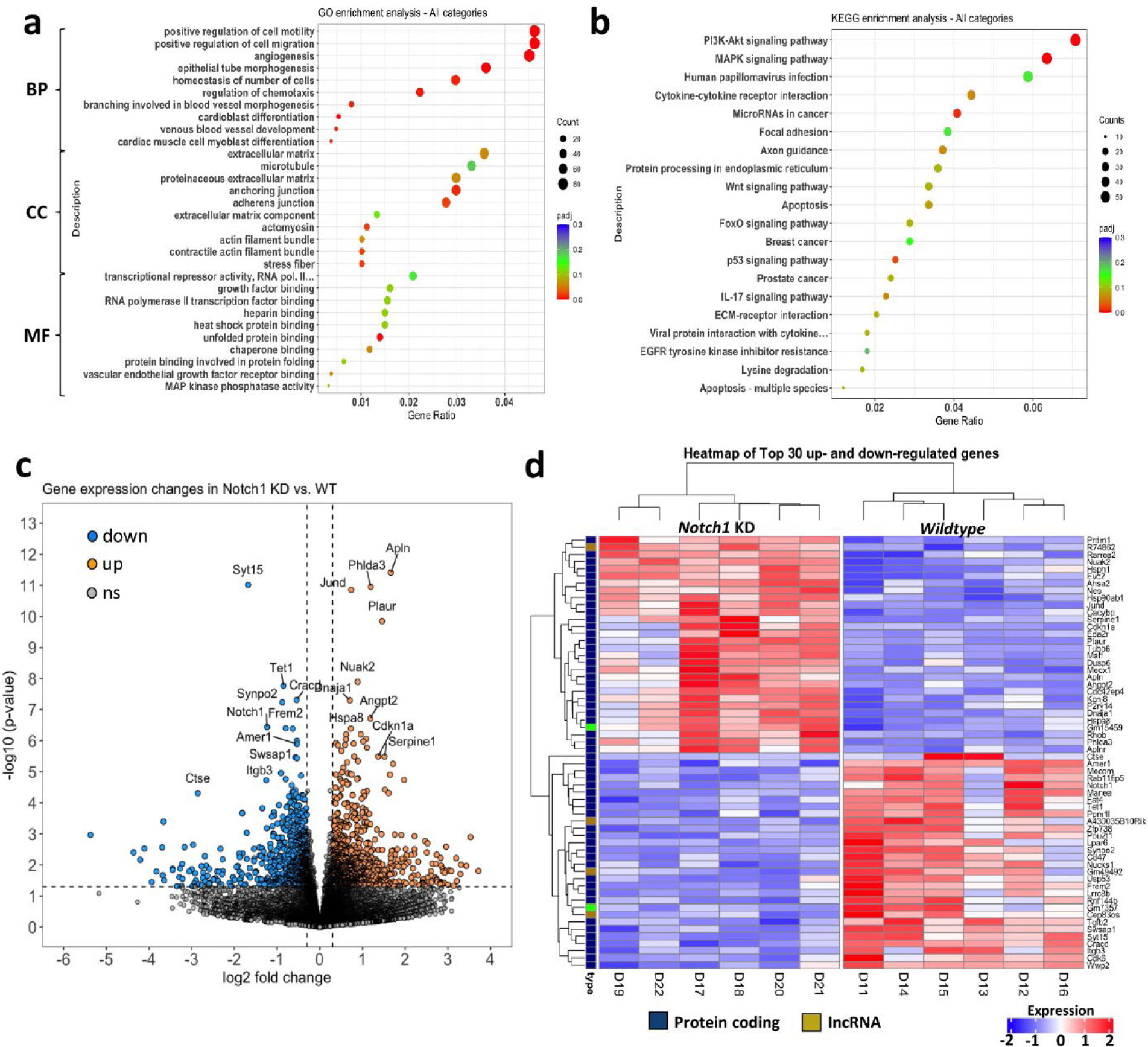
Transcriptional changes in brain endothelial cells following viral knockdown of Notch1. All graphs compare endothelial specific knockdown of Notch1 (n=6 mice infected with AAV-BR1-iCre) to wildtype control endothelial cells (n=6 mice infected with AAV-BR1-eGFP). (**a**) Dot plot for gene enrichment analysis shows top 10 GO terms (for genes up and down regulated) in each GO category after Notch1 knockdown. BP: biological processes, CC: cellular component, MF: molecular function. (**b**) Dot plot shows KEGG classification for differentially expressed genes (top 20 most significantly affected functions) following Notch1 knockdown. (**c**) Volcano plot depicts gene expression based on log2 fold change and statistical significance (adjusted p<0.05). Gene names are attached to the top 10 genes most significantly up (orange) or down (blue) regulated. (**d**) Heat map shows top 30 differentially expressed genes in endothelial cells across each mouse in Notch 1 knockdown or wildtype control groups.

## Discussion

Given there are discrepancies in the literature about basal rates of vascular remodelling, combined with the fact that most studies focus on one or two brain regions, we longitudinally imaged vascular networks across multiple cortical regions. One of our primary findings was that rates of angiogenesis were significantly higher in visual cortex relative to medial/anterior regions such as retrosplenial, motor and FL/HL somatosensory cortex. By contrast, rates of vessel pruning were quite similar across regions. While this is to our knowledge, the first longitudinal imaging study of vasculature in visual cortex, our finding agrees with previous work showing that angiogenesis is quite rare in somatosensory and motor regions^20,31,32^. Our data also indicate that elevated angiogenesis in visual cortex could not be attributed to potential damage related artifacts associated with the cranial window (ie. proximity to the edge of the cranial window) or a higher proportion of ascending venules where angiogenesis typically occurs.

There are several plausible explanations for regional differences in angiogenesis that could work co-operatively or independent of the gene expression differences we observed in the present study. Although speculative, visual cortical areas may have different metabolic and perfusion demands than in medial or anterior cortical regions, which could affect the necessity for producing new vessels over time. Quantitative anatomical studies have shown prominent regional differences in cell density and vascular structure across the dorsal cortex^36,38,40,49,50^. For example, vessel length/branch density and fluid conductance tend to be lower in visual areas relative to retrosplenial, motor and somatosensory cortex. Furthermore, there are significant differences in the density of cell types, such that visual cortex has the highest density of excitatory Glut1 expressing neurons whereas retrosplenial and somatosensory regions are more enriched with inhibitory parvalbumin expressing neurons^40^. Based on our current mapping of angiogenic events across the dorsal cortex (see Fig. 3E), we cannot say with certainty that elevated angiogenesis rates are exclusive to visual cortex since rates were slightly elevated in nearby trunk/whisker somatosensory cortex. Indeed, Kleinfeld’s group has mapped vascular beds across cortex and found they tend to follow regional gradients rather than respect very sharp neuronal or functional borders that typically define cortical areas^51,52^. Thus, a conservative interpretation of our data could be that angiogenesis follows a regional pattern where rates are higher in posterior/lateral regions (ie. Visual and whisker/trunk somatosensory cortex within our imaging window) and lower in medial/anterior cortex. A simple explanation for this is that territories supplied by the posterior, medial or anterior cerebral arteries (CA) possess intrinsic differences in angiogenic potential. Indirect support of this idea comes from recent work showing that vessel loss with aging is generally lower in cortical regions perfused by the PCA than ACA^38^. Further, the PCA has stronger collateral connections than that found in ACA^53^, which could be related to its heightened capacity for angiogenesis. Alternatively, there could be regional differences in “hypoxia pockets” which could provide a stimulus for angiogenesis and have been reported in healthy cortex^54,55^.

Ultimately, the ability for vascular networks to remodel will likely involve changes in gene expression. We therefore leveraged the highly divergent levels of angiogenesis between visual and retrosplenial cortex to search for candidate mechanisms. Since angiogenic signalling mechanisms include co-ordinating interactions between endothelium and nearby pericytes, astrocytes, microglia and neurons, we conducted our original qPCR directed screen in whole tissue (rather than sorted cells) on a select set of genes previously implicated in angiogenesis. Doing so revealed that Notch1 related signalling genes (*Dll4, Jag1, Notch1*) were enriched in retrosplenial cortex, relative to the visual cortex. Using a viral approach for endothelial specific knockdown of *Notch1*, we found that rates of angiogenesis could be significantly increased in both visual and retrosplenial cortex. The fact that angiogenesis could be upregulated in retrosplenial cortex, an area virtually devoid of angiogenesis normally, suggest that Notch1 may normally serve as a brake on this form of plasticity. Notch and VEGF signalling play co-ordinated roles in normal development, as well as diabetes and tumor related angiogenesis that involves spatially/temporally precise patterns of activation^14,56,57^. The present findings agree with previous work showing Notch1 acts to stabilize vasculature^46^, given that endothelial specific deletion of Notch1 in developing retina stimulates endothelial tip cells^15^. Studies in adult animals are much more scant, although one study showed Notch1 knockdown promotes endothelial cell proliferation and reduces elongation in the aorta, consistent with our data in brain^58^. However, most Notch1 knockdown studies in adulthood have been associated with imapirments in ischemia induced angiogenesis in hindlimb and brain^59,60^. On the surface, these previous findings might appear to contradict our findings of increased angiogenesis with Notch1 knockdown. However, in both previous studies, Notch1 required VEGF signalling for the angiogenic response to ischemia. In our study, we did not induce any ischemia (and presumably VEGF signalling) to observe angiogenesis in the healthy brain, nor did it appear that endothelial specific knockdown of *Vegfr2* had a significant effect, although there was a slight trend in visual cortex towards reduced angiogenesis. While the absence of a *Vegfr2* effect may seem surprising since it is essential for vascular development^61^, it is consistent with a recent paper showing no effect of endothelial *Vegfr2* deletion on vascular density in retina, heart or brain^62^. However we do concede a “floor effect” may have been at play since rates of angiogenesis were very low to begin with. Collectively these findings suggest that Notch1 plays a critical role in regulating non-hypoxia/ischemia forms of angiogenesis in the mature mouse cerebral cortex.

Consistent with the idea that VEGF signaling is not essential for Notch1 to regulate adult angiogenesis, our analysis of DEGs in brain endothelial cells after Notch1 knockdown did not reveal significant changes in VEGF isoform or receptor (eg. *Vegfa, Vegfr1 or Vegfr2*) expression. Instead, we found that other angiogenesis related genes such as *Apln, Cdkn1a* and *Angpt2* were some of the most significantly upregulated genes in this group, which is consistent with studies examining Notch1 deletion or inhibition^63,64^. For example, recent work in Zebrafish embryos demonstrated that Apelin was critical for induction of aortic angiogenesis^64^. In agreement with the present work, inhibition of the Notch1 ligand DLL4 stimulated downstream Apelin expression which promoted endothelial sprouting and angiogenesis. The role of Cdkn1a in angiogenesis is not entirely clear as global deletion does not cause developmental defects^65^, but does lead to tumors in adult mice if combined with Cdkn1b deletion^66^. Angiopoietin-2 is a growth factor that has pro-angiogenic properties. Studies have shown that Angiopoietin-2 can stimulate migration and sprouting angiogenesis in endothelial cells with low Tie2 expression^67^, and blocking Angiopoietin-2 can reduce angiogenesis in certain types of cancer such as glioblastoma^68^. Lastly, Notch1 knockdown upregulated *Plaur* and *Serpine1*, that while classically associated with hemostasis, have been implicated in angiogenesis under pathological conditions^69,70^. Given that many of these genes ultimately signal through PI3K/AKT and MAP kinase, it is not surprising to notice these were also strongly affected in the KEGG functional gene analysis.

There are limitations to our study that should be carefully considered. First, we focused our long-term imaging experiments on easily accessible dorsal cortex and therefore omitted regions that extended laterally beyond the parietal bone ridge, such as secondary somatosensory, auditory and insular cortices. Given the fact that these regions would require imaging at a considerable angle and necessitate removal of the mastication related temporalis musculature, future imaging studies might be better served with an implanted two-photon endoscope/fiberscopes^71^. Furthermore, the considerable amount of time required to collect high-resolution images of cortical vasculature, prompted us to restrict our imaging to 2-4 cortical regions and a depth of 400µm below the cortical surface per mouse. Thus our estimates of angiogenesis and pruning do not include deeper cortical regions involving layers 5 and 6. Since there are layer specific differences in cerebral blood flow, vessel length and cell density, it is conceivable that rates of angiogenesis and pruning may differ in those deeper layers^40,72^. It is worth noting that our result showing that angiogenesis varies as a function of superficial cortical depth (0-400µm, see Fig. 4c), closely aligns with previous imaging work where hypoxia induced capillary sprouting was most abundant within 250µm from the cortical surface^33^. A final caveat that deserves mention is we focused most of our experiments on C57BL/6 mice. Strain differences in vascular volume and length have been reported in the literature, as wella as differences in the angiogenic response to hypoxia^51,73^. Indeed we did find that basal rates of angiogenesis tended to be lower in our *Vegfr2* knockdown experiments which involved mice on a CD1 background. Thus there could have been a floor effect that occluded our ability to detect a decrease in angiogenesis. Since many mouse strains are commonly used in research, future studies would be required to specifically resolve strain related differences in angiogenesis.

What is the functional significance of our findings? We know that angiogenesis is critical for proper brain development, but also plays a contributing role in pathology associated with neurological conditions and correlates with recovery from different forms of injury in the adult brain^21,23,26,74,75^. Even in the absence of injury or disease, some studies suggest that angiogenesis may participate in every day activities like learning and memory or the rapid cognitive boosting effects of exercise^29,76,77^. While we did not enrich our socially/group housed mice with exercise wheels or learning tasks, the production of new vessels, even when stimulated with Notch1 knockdown, usually took many days or weeks to occur. Further, if one assumes there are 1-2×10^4^ capillaries per mm^3^ of cortex, new or pruned vessels would affect less than 1% of the total capillary network over our 3-week imaging period. Therefore, we think that ongoing cerebral angiogenesis and pruning in the normal brain is most relevant to slow changes, such as those associated with aging. Indeed, our group has recently shown that visual cortex resists age related vessel loss whereas retrosplenial, motor and somatosensory limb cortex exhibit significant decline on the order of 10% over 18-22 months^38^. It is therefore conceivable that elevated rates of angiogenesis in visual cortex may engender resiliency to the effects of aging. Whether angiogenesis significantly affects blood flow over the long term (ie. weeks to years) remains to be determined. At the very least, our quantitative data could help inform future computational studies that model how structural plasticity of vessels augments blood flow over time.

## Methods

### Animals

Two to four-month old male and female C57BL/6J mice were used for cortical imaging experiments. For endothelial specific knockdown experiments, we used adult mice homozygous for the floxed *Vegfr2/Flk1* gene with a CD1 background (generous gift from Dr. Jane Rossant), or mice homozygous for the floxed *Notch1* gene (N1CKO, JAX# 007181) backcrossed to a C57BL/6J background^78^. All animals involved in this study were cared for and treated following protocols approved by the University of Victoria Animal Care Committee and are in compliance with the guidelines set by the Canadian Council on Animal Care (CCAC) standards. The animals were housed in standard cages with ad libitum access to water and the standardized laboratory diet. Mice were kept in a room maintained at 22.5°C ±2.5°C in a 12-hour light/dark cycle and a humidity-controlled environment. Reporting of this work complies with ARRIVE guidelines.

### AAV based manipulation of endothelial Notch1, Vegfr2 or Tdtomato reporter expression

AAV-BR1-iCRE or eGFP was prepared as previously described^48^. For determining the efficacy and specificity of endothelial expression, three *Gt(ROSA)26Sor^tm9(CAG-tdTomato)Hze^* with a C57BL6/J background (“Ai9” reporter mice) were intravenously injected with AAV-BR1-iCRE (20µl of 5.0×10^12^ GC/mL mixed with 80µl of 0.9% saline). Endothelial specific knockdown of *Vegfr2* or *Notch1* was achieved by intravenous injection of AAV-BR1-iCRE (as described above) into homozygous *Notch1* or *Vegfr2* floxed mice. A subset of mice received intravenous injection of AAV-BR1-eGFP control virus. Following injection of virus, mice were given a 9-10 day rest period before imaging to allow transfection of endothelial cells.

### Cranial window preparation

To implant cranial windows for longitudinal imaging, mice were anesthetized throughout the surgical procedure using isoflurane gas with medical grade air (80% N2, 20% O2) at a flow rate of 0.7L/min. The isoflurane vaporizer was set to 2% for induction until the animal reached an anesthetic depth appropriate for transfer to the surgical stage and the vaporizer was set at 1.3% for maintenance throughout the surgery. Mice were head fixed to a custom-built surgical stage and a feedback driven heating pad was used to maintain body temperature between 36-37°C. Mice received an injection of 30µL of 2% dexamethasone (i.p. Vetoquinox; Dexamethasone sodium phosphate) to reduce surgery related inflammation and lidocaine (s.c. 30µL at 20mg/ml) beneath the scalp as an analgesic. A mid-line incision was done to expose the skull surface which was cleaned and dried with a sterile cotton swab. For posterior cranial windows, a custom metal ring (11.3 mm outer diameter, 7.0 mm inner diameter, 1.5mm height) was secured to the skull with dental cement over the right hemisphere and above the retrosplenial, visual and somatosensory cortices, based on stereotaxic co-ordinates. For anterior cranial windows, the metal ring was secured to the skull above motor cortex and forelimb somatosensory areas. A 4mm diameter craniectomy was made by carefully thinning a circular area of the skull with a high speed dental drill. Cold HEPES buffered artificial cerebral spinal fluid (ACSF) was regularly applied to the skull to prevent heating. Once the skull was thin enough to visualize underlying vessels, the skull was removed with sterile forceps while leaving the dura intact. Gel foam soaked in ACSF was used to control for any bleeds that occurred throughout the surgery. A 5mm diameter circular glass coverslip was positioned over the craniotomy, secured with cyanoacrylate glue and dental acrylic around the circumference of the coverslip. Mice were monitored in the acute post-surgery period as they recovered under a heat lamp or pad. If recovery was normal, mice were returned to their home cage where they were monitored regularly for 4 weeks prior to beginning the longitudinal 2-photon imaging timeline. Mice who did not have a clear window 4 weeks post-surgery were excluded from imaging experiments.

### Intrinsic optical signal imaging

Intrinsic optical signal (IOS) imaging was used to identify cortical sensory regions by measuring changes in deoxygenated hemoglobin in response to sensory stimuli. Mice with a clear cranial window (no damage to the dura, extensive bone growth or inflammation) were anesthetized with isoflurane mixed with medical grade air (2% isoflurane for induction, 1.0% for maintenance) and transferred to a custom-built head fixing stage. Body temperature was monitored and maintained at 36-37°C using a rectal probe and heating pad. The brain surface was imaged with a 2X objective (NA=0.14) on an upright Olympus microscope connected to a MiCAM02 CCD camera and BrainVision software (SciMedia). An image of the surface blood vessels was captured and then a red LED (625nm) was used to illuminate the cortical surface. Either the left eye, whisker, forelimb (FL) or hindlimb (HL) was stimulated to evoke responses within their respective cortex. For tactile stimulation, a pencil lead attached to a piezo-electric wafer was used to stimulate the FL or HL using 5ms biphasic pulses at 100Hz for 1s per trial. A single whisker on the contralateral face was stimulated with a small loop attached to a pencil lead using 3ms biphasic pulses at 25Hz. Visual cortical responses were elicited by illuminating the left eye using a cyan LED (505nm). Each set of stimulation trials was composed of 10-12 individual stimulation trials that was followed by a no-stimulus trial and a 12s interval between stimulation/no stimulation trials. Within each trial, 3s of reflected red light was collected with 1s of sensory stimulation starting after 1s of baseline. Each 3s trial was imaged at 100 Hz with a 10ms exposure per frame.

For processing IOS images in ImageJ, the 10-12 Stim/no stim trials were averaged together and mean filtered using a 5-pixel radius. The baseline surface reflectance image (Ro) was generated by averaging the first 100 pre-stimulus image frames. The baseline image (Ro) was than subtracted from the 300 frames stim/no stim averaged frames and then divided to generate a dR/Ro. To clearly identify each cortical region, we mean projected 150 frames starting 0.5s to 1.5s after stimulation when the initial change in deoxyhemoglobin signals is maximal. The image was then thresholded at 70% of peak response values, the borders of each functional area were demarcated and superimposed onto a cortical image with visible superficial vasculature.

### *In vivo* 2-photon imaging and analysis

Approximately 5 weeks after cranial window surgeries, mice with a clear window were anesthetized using isoflurane mixed with medical grade air (2% Isoflurane for induction, 1% maintenance). Once appropriate anesthetic depth was reached, mice were transferred to a custom-built stage which fixed the head holding ring into place. Body temperature was monitored and maintained at 36-37°C. C57BL/6J mice were imaged longitudinally on days 0, 2, 9-10, and 23. These time intervals were based on previous imaging studies describing vascular remodelling associated with micro-emboli^31^. Genetic knockdown experiments followed a similar timeline except we omitted day 2 which we deemed unnecessary given the increased total number of imaging sessions. Prior to the start of each imaging session, mice were intravenously injected with FITC-dextran (100µL of 3-5% w/v in 0.9% saline; 70kDa, Sigma #46945) or Texas Red dextran (100µL of 3% w/v in 0.9% saline; ThermoFisher; MW 70kDa, D1830) to fluorescently label blood plasma. High-resolution *in vivo* images of the cerebral vasculature were acquired using a multiphoton laser scanning microscope running Fluoview FV10-ASW software (Olympus FV1000MPE) and a mode-locked Ti:Sapphire laser (Spectral Physics) with a water dipping 20X objective lens (Olympus XLUPlanFl, NA=0.95). The laser was tuned to 800nm for FITC dextran or 850nm for Texas Red dextran. Laser power was adjusted according to imaging depth and ranged between 15mW to 150mW from superficial to deep images. Emitted light was split by a 570nm dichroic mirror prior to passing through bandpass emission filters (495-540nm and 575-630nm). Imaging specifications were as follows: z-step of 2µm, dwell time of 2µs/pixel up to a depth of 420µm and with an imaging area of 1024×1024 pixels (635×635µm^2^ across x-y planes). For each mouse, 2-4 brain regions were imaged using IOS generated maps for visual, FL, HL or whisker cortex, as a reference. Retrosplenial and motor cortex were demarcated based on their relative position to the midline and V1 or FL/HL somatosensory areas, respectively. Imaging of the same areas over time was achieved by collecting brightfield images of surface cortical vessels which provided landmarks from week to week.

To quantify rates of angiogenesis and pruning across different experimental conditions, image stacks for the first and last imaging time-point (eg. days 0 vs. 23 or day 33 vs. 56) were opened and aligned using Fiji software (ImageJ 1.53q). Each image stack was binned into maximum intensity z-projections that each consisted of 20-30 image frames. Z-projection images between the two time-points were compared by an observer blind to condition. Suspected angiogenic or pruning events were further confirmed by manually inspecting original 3D image stacks and examining intermediate time-points (eg. day 9 or 42). An angiogenic event was recorded if a fully formed and connected vessel was present on D23 that was either completely absent or accompanied by a sprout at D0. Also included were rarer cases where a sprout (not connected to another vessel) was evident at D23 that was preceded by no sprout (“absent”) or a shorter sprout on D0, that grew at least 10µm in length. Pruning events were defined as when a vessel or sprout that was present on D0, was completely absent on D23. The rate of turnover was calculated by summing the number of angiogenic and pruning events in each region, divided by the total volume sampled.

For plotting the density of angiogenic and pruning events for all imaging stacks relative to bregma (see Fig. 3E), we first used IOS maps of the hindlimb region as a reference landmark (common to both anterior and posterior cranial windows, assume centroid is 1.5mm lateral and 0.5mm posterior to bregma) for establishing stereotaxic co-ordinates of all other imaging stacks. Each angiogenic or pruning event was then plotted in 2D space in Python 3.9.12 (https://www.python.org/downloads/release/python-3912/) and then interpolated with a Kernel Density Estimate using the Seaborn package (https://seaborn.pydata.org/generated/seaborn.kdeplot.html) in a Juypter notebook (https://jupyter.org/). The Seaborn KDE plot forms a continuous probability density curve of the (x,y) data in 2D space using a bandwidth method of 0.05. Color scale is the proportional density for each plot, from maximum to minimum values in 10 level increments. Total area under all densities was normalized to sum to 1.

### Histology and confocal imaging

Adult or 15 day old mice were euthanized with an overdose of sodium pentobarbital (i.p. 50µL of Euthanyl at 120mg/mL, Bimedia-MTC Animal Health Inc.). Once toe-pinch reflexes were lost, mice were transcardially perfused with 10mL of 0.1M phosphate buffered saline (PBS). Following dissection, brains was immersed in 4% paraformaldehyde (PFA) overnight and later transferred to a 30% sucrose solution (in 0.1M PBS with 0.2% sodium azide). Brains were sectioned on a freezing microtome (American Optical Corp.) into 40-50µm thick coronal sections and stored in a 12-well plate with 0.1M PBS in 0.2% sodium azide. For immunostaining, sections were incubated overnight in 0.1M PBS + 0.1% TX-100 solution containing primary antibody for rat anti-CD31 (label endothelial cells; 1:200 dilution, BD Clone MEC 13:3: #553370) or sheep anti-CD93 (label endothelial tip cells, 1:500 dilution, R&D Systems, AF1696). After 3 washes, sections were incubated in PBS containing Cy5 or Alexa 488 conjugated secondary antibodies (1:400) at room temperature for 4 hours. Sections were washed, mounted onto gelatin coated slides, allowed to dry and cover-slipped with Fluoromount-G (Southern Biotech).

Confocal image stacks were collected under a 10 or 20X objective lens (NA= 0.4 or 0.75, respectively) using Fluoview FV10-ASW software (Olympus Corp.). A 488, 561 or 635nm laser line with 4µs pixel dwell time was used to excite tdTomato or fluorophore conjugated secondary antibodies. Kalman averaged images (Kalman=2 frames) were collected at 4µm z-steps covering 1271×1271µm (1.24µm/pixel) under 10X magnification, or 2µm z-steps covering 424×424µm (0.26µm/pixel) under 20X magnification. To assess the efficacy of the AAV-BR1-driven Cre-recombinase, tdTomato signal was overlaid against CD31 immunolabeling of endothelial cells. To do this, images from each channel were first maximally projected and de-noised with median filter (radius=0.8 pixels). Background fluorescence was subtracted from each image (by median filtering original image with 50 pixel radius). The resultant background corrected imaged were binarized using a Li threshold and then overlaid on top of each other. Pixel overlap for each channel was quantified and expressed as a percentage using Fiji software (ImageJ 1.53q).

### Whole tissue RNA extraction and RT-qPCR

Mice was overdosed by injection of sodium pentobarbital (i.p. injection of 240mg/mL) and transcardially perfused with 10mL of 0.1M phosphate buffered saline to remove the blood. Brain tissue was quickly extracted, and sliced into 1mm thick coronal sections. The retrosplenial and visual cortex were micro-dissected and flash-frozen in liquid nitrogen. Total RNA was extracted from frozen tissue using RNeasy Mini kit (Qiagen, cat#74104) with additional DNase treatment (Qiagen, cat#79254). 100ng of RNA per sample was used to prepare cDNA using High-Capacity cDNA synthesis kit (Applied Biosystems, cat#4368814). cDNA was then diluted 5-fold for RT-qPCR runs. Primers (see below) were designed using NCBI portal and Primer Bank to detect the levels of the indicated transcripts and those selected achieved an efficiency rate between 90-110%. To further validate the selection of primers, specificity was calculated first using NCBI online primer blast tool in conjunction with UCSC genome browser and later confirmed experimentally using 10-fold serial dilutions and melt curve analyses ran in triplicates. Fluorescent signals were acquired using the StepOne plus system with qPCR reaction mixtures comprised of: 1µL cDNA, 0.5µL of each primer, 3µL RNase, DNase free water, and 5µL of SYBR Green Master Mix (Applied Biosystems). Thermocycling conditions for reactions were as follows: 50°C for 2 minutes, 95°C for 2 minutes followed by 40 cycles of denaturation at 95°C for 15 seconds and annealing at 60°C/62°C for 1 minute. Triplicate reactions were performed for each sample. To analyse RT-PCR results delta-delta Ct methods were used, where Ct values were averaged and normalized to the expression of geometric mean of housekeeping genes *Tbp,* and *Hprt* to calculate relative levels of mRNA expression of genes of interest using Design and Analysis Software Version 2.4.3 (Applied Biosystems). Values for each gene in retrosplenial cortex were then normalized to expression in visual cortex. Forward and reverse primers used for RT-qPCR were as follows: *Tbp* (housekeeping): CCCCACAACTCTTCCATTCT and GCAGGAGTGATAGGGGTCAT; *Hprt* (housekeeping): AGCCTAAGATGAGCGCAAGT and TTACTAGGCAGATGGCCACA; *Evl*: CCGTGATGGTCTACGATGACA and GTCCCCGGCAGTTGATGAG; *Adam17:* AGGACGTAATTGAGCGATTTTGG and TGTTATCTGCCAGAAACTTCCC; *Dll4*: ATGGTGGCAGCTGTAAGGACC and AGGCATAACTGGACCCCTGG; *Notch1*: CCCTTGCTCTGCCTAACGC and GGAGTCCTGGCATCGTTGG; *Vegfr1 (Flt1)*: TGGCTCTACGACCTTAGACTG and CAGGTTTGACTTGTCTGAGGTT*; Vegfr2 (Flk1)*: TTTGGCAAATACAACCCTTCAGA and GCAGAAGATACTGTCACCACC; *Jag1*: CCTCGGGTCAGTTTGAGCTG and CCTTGAGGCACACTTTGAAGTA.

### Endothelial cell separation

Mice were deeply anesthetized with 2% isoflurane and euthanized by decapitation. Brain was extracted and immediately transferred to sterile 35mm petri dish containing 1mL HBSS with no calcium and magnesium (Thermo Fisher #14175095). Brain tissues were cut into small pieces using scalpel before transferring them into a 15mL falcon tube. The mechanical dissociation of brain tissue was performed using the Adult Brain Dissociation Kit (Miltenyi Biotec #130-107-677) as per manufacturer’s instructions. Briefly, the brain was homogenized by gently pipetting up and down ∼10 times with a 1mL pipette at 37⁰C. The recovered homogeneous cell mixture was gently applied to a smart strainer (Miltenyi Biotec #130-098-462) to remove the connective tissue. In the following steps, samples were always kept on ice unless otherwise indicated. Myelin was removed from the cell mixture using Myelin Removal Kit (Miltenyi Biotec #130-096-733) as per manufacturer’s instructions and then passed through LS columns (Miltenyi Biotec #130-042-401) using Quadro-MACS separator (Miltenyi Biotec #130-090-976). After 3 washes with Auto-MACS rinsing solution (Miltenyi Biotec #130-091-222) containing 0.5% BSA (Miltenyi Biotech #130-091-376), the flow through was centrifuged for 10min and re-suspended in Auto-MACS Rinsing Solution containing 0.5% BSA. CD45 positive cells were removed from the total cell suspension using CD45 microbeads (Miltenyi Biotec #130-052-301) by passing the cell mixture through LS columns. The flow through containing CD45-negative cells was then centrifuged for 10min. The cell pellet was re-suspended in Auto-MACS Rinsing Solution with 0.5% BSA and incubated with CD31 microbeads (Miltenyi Biotech #130-097-418) and passed through MS column (Miltenyi Biotech #130-042-201) using Octo-MACS separator (Miltenyi Biotec #130-042-108). After 3 washes with Auto-MACS rinsing solution with 0.5% BSA, column bound CD31 positive endothelial cells were collected by adding 1mL Auto-MACS rinsing solution with 0.5% BSA. The collected sample volume was centrifuged at max speed in a benchtop centrifuge for 10 minutes and after discarding the supernatant, the endothelial cell pellet was flash frozen and kept at −80⁰C until further use.

### RNA sequencing and bioinformatics

RNA sequencing was performed by Novogene Inc. according to their standardized procedures. For library construction, messenger RNA was purified from total RNA using poly-T oligo-attached magnetic beads. After fragmentation, the first strand cDNA was synthesized using random hexamer primers, followed by the second strand cDNA synthesis using either dUTP for directional library or dTTP for non-directional library^79^. For the non-directional library, samples were ready after end repair, A-tailing, adapter ligation, size selection, amplification, and purification. For the directional library, samples were ready after end repair, A-tailing, adapter ligation, size selection, USER enzyme digestion, amplification, and purification. The library was checked with Qubit and real-time PCR for quantification and bioanalyzer for size distribution detection. Quantified libraries were pooled and sequenced on Illumina platforms, according to effective library concentration and data amount.

For data quality control, raw data (raw reads) of fast q format were firstly processed through in-house perl scripts. In this step, clean data (clean reads) were obtained by removing reads containing adapter, reads containing ploy-N and low quality reads from raw data. At the same time, Q20, Q30 and GC content the clean data were calculated. All the downstream analyses were based on the clean data with high quality. Reference genome and gene model annotation files were downloaded from genome website directly. Index of the reference genome was built using Hisat2 v2.0.5 and paired-end clean 1 reads were aligned to the reference genome using Hisat2 v2.0.5. We selected Hisat2 as the mapping tool for that Hisat2 can generate a database of splice junctions based on the gene model annotation file and thus a better mapping result than other non-splice mapping tools^80^.

Quantification of gene expression was done using featureCounts v1.5.0-p3 to count the read numbers mapped to each gene^81^. The expected number of Fragments Per Kilobase of transcript sequence per Millions base pairs (FPKM) of each gene was calculated based on the length of the gene and read counts mapped to a particular gene. FPKM considers the effect of sequencing depth and gene length for the read count at the same time, and is currently the most commonly used method for estimating gene expression levels. Differential expression analysis of the two groups was performed using the DESeq2R package (1.20.0). DESeq2 provide statistical routines for determining differential expression in digital gene expression data using a model based on the negative binomial distribution. The resulting P-values were adjusted using the Benjamini and Hochberg’s approach for controlling the false discovery rate. Genes with an adjusted P value ≤ 0.05 found by DESeq2 were assigned as differentially expressed. Corrected P value ≤ 0.05 and a Log2 fold change of 1.3 were set as the threshold for significant differential expression. Gene Ontology (GO) enrichment analysis of differentially expressed genes was implemented by the cluster Profiler R package, in which gene length bias was corrected. GO terms with corrected P value ≤ 0.05 were considered significantly enriched by differential expressed genes. We used cluster Profiler R package to test the statistical enrichment of differentially expressed genes in KEGG pathways^82^.

### Statistics

All data were statistically analysed using GraphPad Prism (versions 7 and 8, RRID:SCR_002798). Statistical tests for each experiment are reported in the Figure legends. To analyze between group differences (ie. when only 2 groups compared on a single factor), planned two-tailed student’s *t* tests (paired or unpaired) were employed as appropriate. One-way ANOVA was used to compare region specific differences whereas two-way ANOVAs were used to compare differences related to factors such as sex and brain region. Significant main effects from ANOVAs were analyzed with Tukey’s multiple comparisons tests. Linear regression tests were used to determine if there was a relationship between: a) the quantity of angiogenic and pruning events within the same region and b) an imaging area’s proximity to the edge of the cranial window and rates of angiogenesis. A two-tailed Mann-Whitney test was used to compare differences between arteriole and venous branch order distributions with respect to formation or elimination of capillaries. *P*<0.05 was considered statistically significant.

## Data Availability

Data generated for this study are available from the corresponding author on reasonable request.

## Acknowledgements

We are grateful Roobina Boghozian, Ana Paula Cota for their assistance with experiments, as well as Angie Hentze and Taimei Yang for managing the mouse colony. We thank Animal Care Services for their help with animal husbandry and welfare. This research was supported by operating, salary and equipment grants to C.E.B. from the Canadian Institutes of Health Research (CIHR), Heart and Stroke Foundation (HSF), Natural Sciences and Engineering Research Council (NSERC).

## Author Contributions

C.E.B, A.R., B.S. conceived the study. C.E.B, A.R wrote the manuscript. J.K provided critical reagents, A.R., B.S., D.H., S.S., K.N., P.R., M.C., and C.E.B. performed experiments and/or analyzed data.

