## Supplementary data for "Angiogenesis in the mature mouse cortex is governed in a region specific and Notch1 dependent manner"

**Supplementary Figure 1**

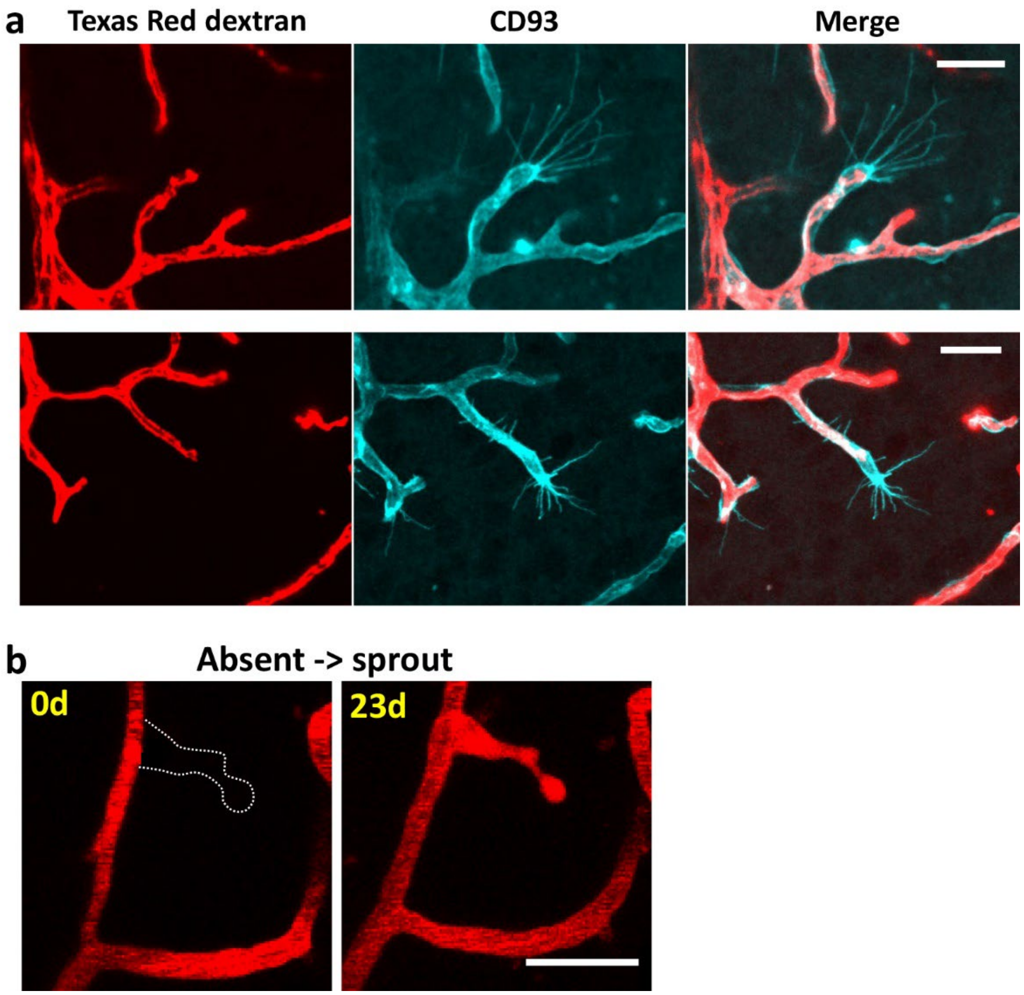


**Supplementary Figure 1. Images show distribution of dye labeled blood plasma in growing blood vessels in visual cortex at post-natal day 15 or in adulthood.** (**a**) Top and bottom rows: confocal images from coronal brain sections show two examples of Texas Red labeled blood plasma inside the lumen of growing endothelial cells immunolabelled with CD93. Note that dye labelled blood plasma fills the lumen of these growing vessels that possess a Tip cell at one end. (**b**) *In vivo* 2-photon images show appearance of vessel sprout. Note that the lumen of the sprout is filled with fluorescently labelled blood plasma. Scale bars = 20µm.

**Supplementary Figure 2.**

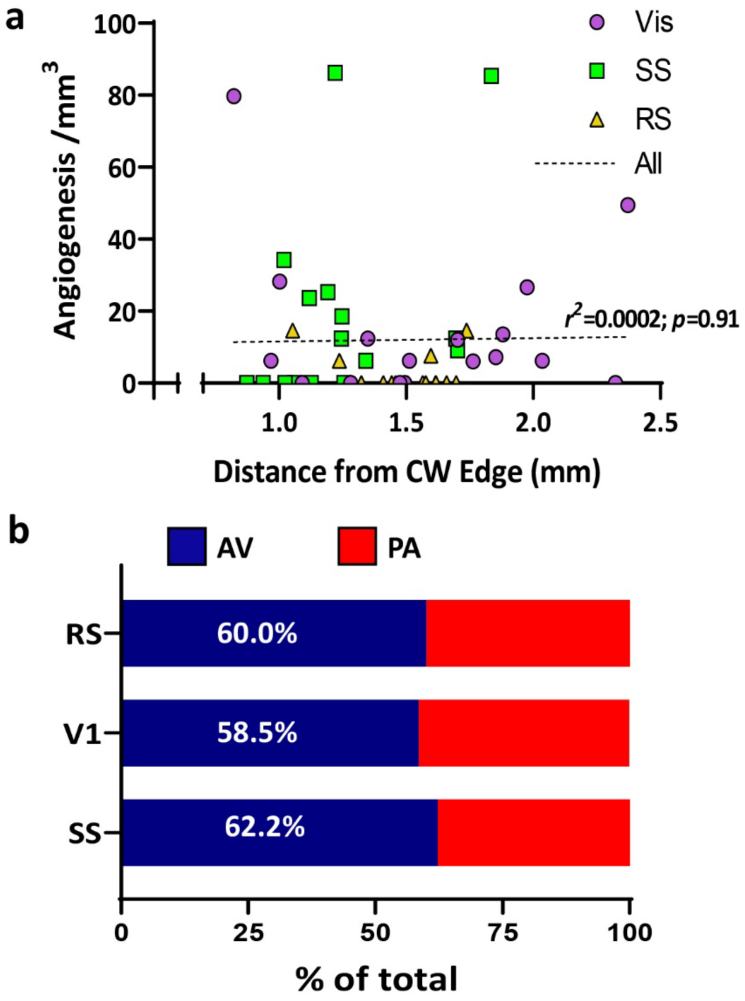


**Supplementary Figure 2. Regional differences in angiogenesis are not attributable to their relative positon to the edge of the cranial window or relative proportion of ascending venules.** (**a**) Linear Regression analysis showing no relationship between the rate of angiogenesis for each cortical region (visual, somatosensory and retrosplenial), and its distance to the nearest edge of the cranial window. (**b**) Graph shows the % of ascending venules (AV) and penetrating arterioles (PA) in each cortical region.

### Supplementary Figure 3

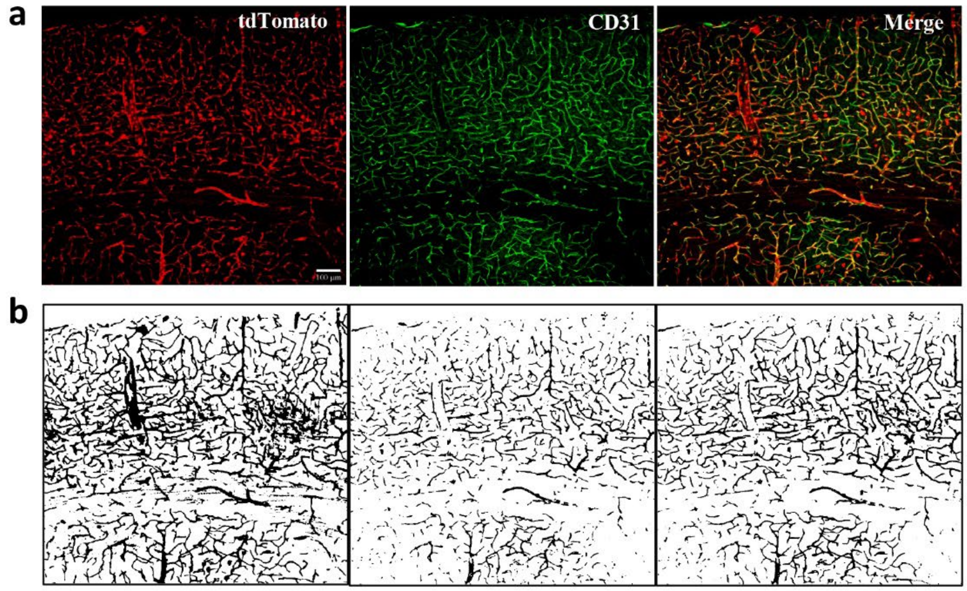


c

| | tdTomato overlap with CD31 | CD31 overlap with tdTomato |
|---|---|---|
| S1 – Barrel Cortex | 89% | 92% |
| S1 – Front Limb | 86% | 89% |
| Retrosplenial Cortex | 86% | 90% |
| Visual Cortex | 84% | 89% |
| **Average** | **86%** | **90%** |

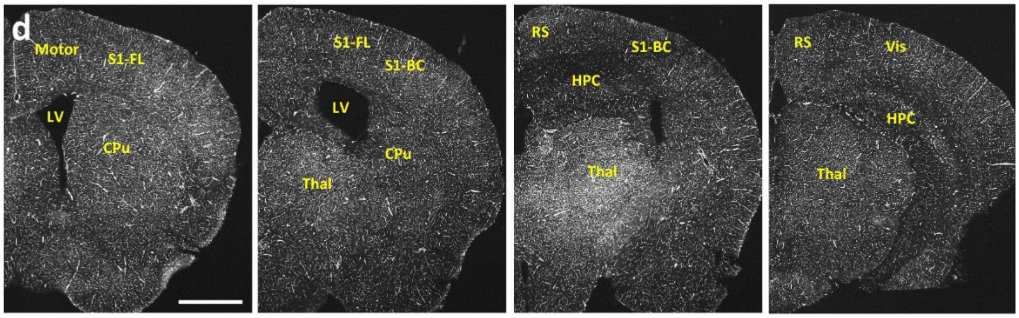


**Supplementary Figure 3. Injection of AAV-BR1-iCRE efficiently transfects most vascular endothelial cells. (a)** Confocal image montage showing tdTomato reporter expression and endothelial specific protein CD31 in reporter mouse (Ai9) intravenously injected with AAV-BR1-iCre. Note the extensive and specific expression of tdTomato in CD31 expressing vessels. (**b**) Shows binarized pixels for each fluorescent image after thresholding. (**c**) Table shows the % of tdTomato pixel overlap with CD31 and CD31 overlap with tdTomato in different cortical regions. (**d**) Montage shows extensive Cre dependent tdTomato reporter expression throughout brain's vasculature. LV: lateral ventricle, CPu: Caudate Putamen; Thal: thalamus; HPC: hippocampus, Vis: visual cortex, S1: primary somatosensory cortex. Scale bar = 100µm in a; 1mm in d.

**Supplementary Figure 4**

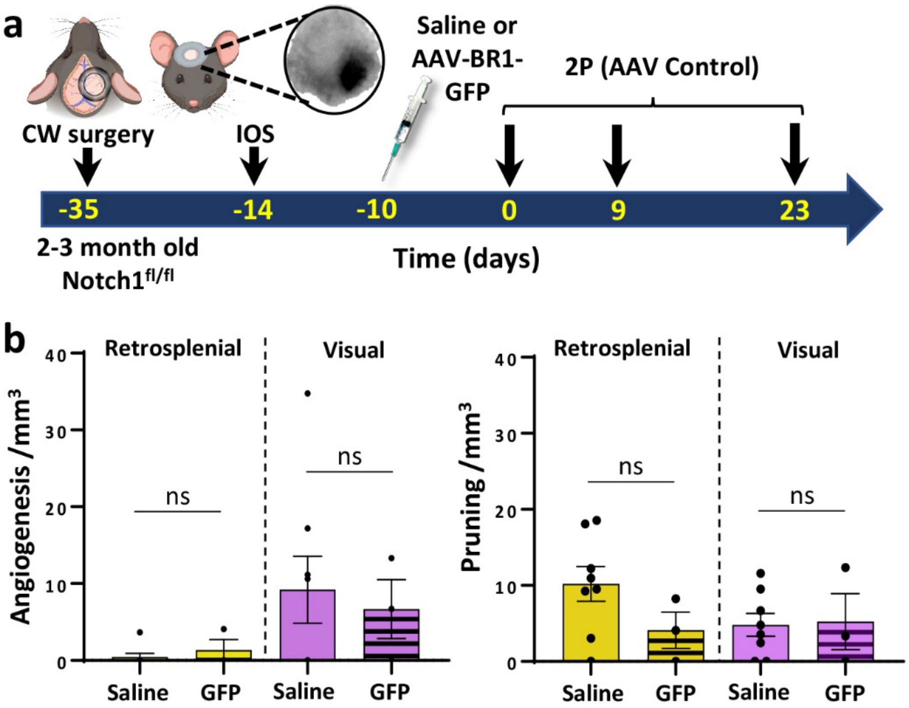


**Supplementary Figure 4. Injection of control AAV-BR1-GFP virus does not alter rates of angiogenesis or pruning.** (**a**) Timeline of experimental procedures and 2-photon (2P) imaging sessions. (**b**) Bar graphs show rate of angiogenesis (left) or pruning (right) in retrosplenial and visual cortex over the 23 day imaging period following injection of saline or AAV-BR1-GFP (n = 8 and 3 mice, respectively). Data in b were analysed with unpaired two-tailed t-tests. ns = not significant. Data are mean ± SEM.

**Supplementary Figure 5**

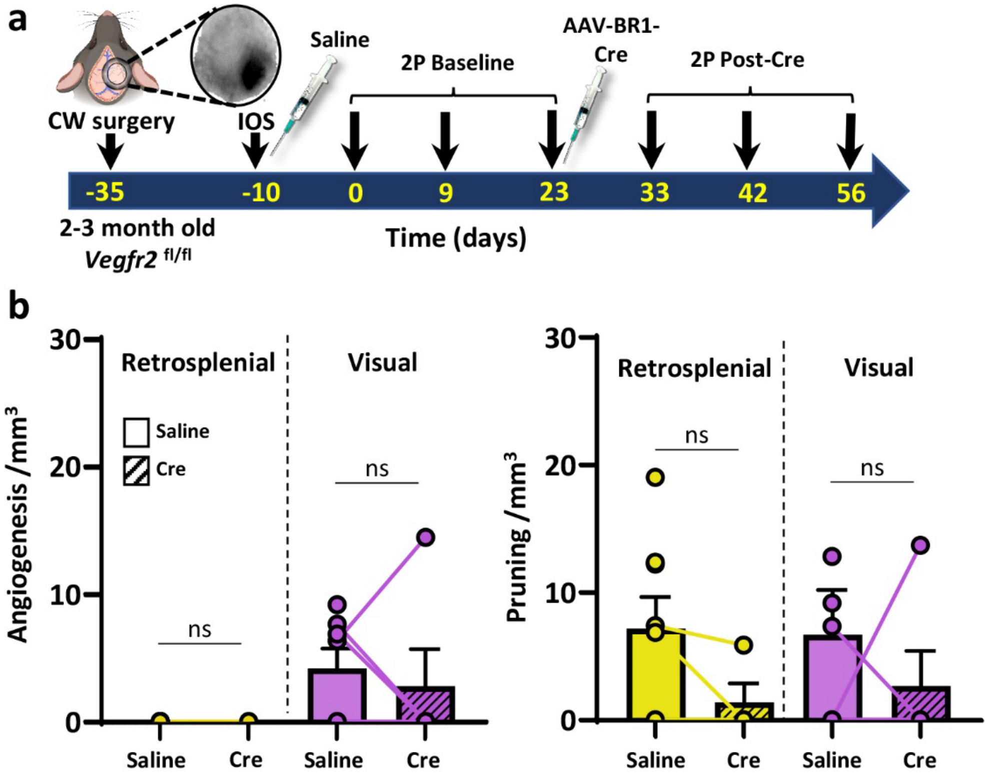


**Supplementary Figure 5. Endothelial specific knockdown of *Vegfr2* does not affect cerebral angiogenesis or pruning.** (**a**) Timeline of experimental procedures and 2-photon (2P) imaging sessions. (**b**) Graphs show rate of angiogenesis (left) or pruning (right) in retrosplenial and visual cortex in *Vegfr2* floxed mice following treatment with saline (baseline) and then AAV-BR1-iCre (n = 8 mice total, with 6 mice completing both baseline and post-AAV-BR1-iCre imaging). Data in b analysed with two-tailed Mann-Whitney tests. ns = not-significant. Error bars: mean ± SEM.

### Supplementary Figure 6

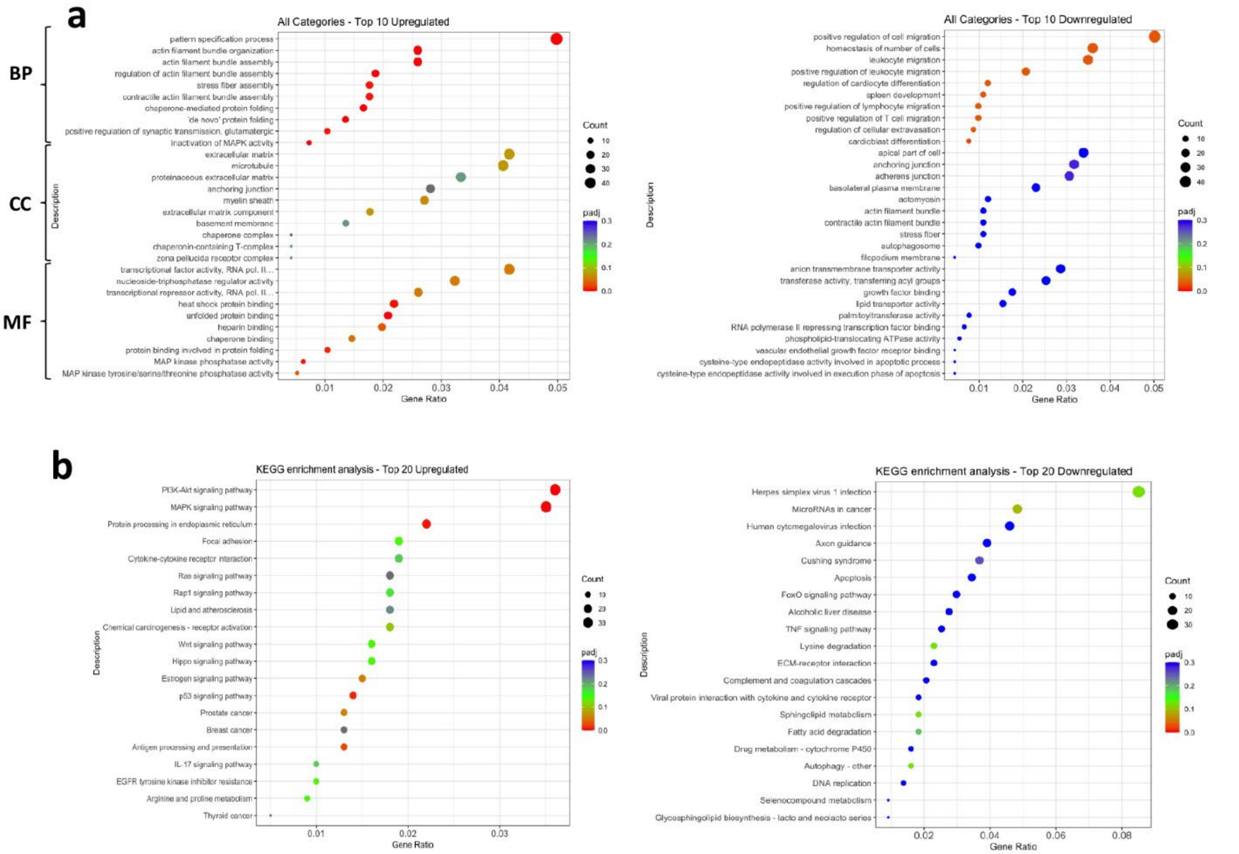


**Supplementary Figure 6.** Dot plots show GO (a) and KEGG (b) analysis comparing top upregulated (left side) or down-regulated (right panel) DEGs in brain endothelial cells isolated from Notch1 knockdown mice compared to wild-type controls (n=6 mice per group).
